# Concurrent emergence of view invariance, sensitivity to critical features, and identity face classification through visual experience: Insights from deep learning algorithms

**DOI:** 10.1101/2024.06.08.597949

**Authors:** Mandy Rosemblaum, Nitzan Guy, Idan Grosbard, Libi Kliger, Naphtali Abudarham, Galit Yovel

## Abstract

Visual experience is known to play a critical role in face recognition. This experience is believed to enable the formation of a view-invariant representation, by learning which features are critical for face identification across views. Discovering these critical features and the type of experience that is needed to uncover them is challenging. We have recently revealed a subset of facial features that are critical for human face recognition. We further revealed that deep convolutional neural networks (DCNNs) that are trained on face classification, but not on object categorization, are sensitive to these facial features, highlighting the importance of experience with faces for the system to reveal these critical features. These findings enable us now to ask what type of experience with faces is required for the network to become sensitive to these human-like critical features and whether it is associated with the formation of a view-invariant representation and face classification performance. To that end, we systematically manipulated the number of within-identity and between-identity face images and examined its effect on the network performance on face classification, view-invariant representation, and sensitivity to human-like critical facial features. Results show that increasing the number of images per identity as well as the number of identities were both required for the simultaneous development of a view-invariant representation, sensitivity to human-like critical features, and successful identity classification. The concurrent emergence of sensitivity to critical features, view invariance and classification performance through experience implies that they depend on similar features. Overall, we show how systematic manipulation of the training diet of DCNNs can shed light on the role of experience on the generation of human-like representations.

---

Object recognition is a computationally challenging task that humans resolve effortlessly. To successfully classify objects into different categories, the brain must create an identity-preserved representation that is tolerant to within-class changes, such as viewpoint, lighting, size, occlusion and so forth (1). This is achieved by emphasizing features that remain unchanged across different variations, while disregarding features that vary across these variations (2). The nature of the experience that is required for the visual system to learn which features are critical and generate a view-invariant representation have so far remained unknown.

Recent advancements in machine vision have successfully resolved the task of object and face recognition with deep convolutional neural networks. These algorithms, trained on thousands of images in a supervised or self-supervised manner, now perform on par with humans in face and object classification (3). Whereas the exact computations employed by these algorithms and their similarity to the computations used by humans to resolve this task are unknown, recent studies have uncovered notable similarities between the representations generated by DCNNs and the human brain and mind (4–7). Thus, by studying the type of experience that is required to generate *human-like* representations in DCNNs, we can gain insights on the ingredients that are needed for these representations to emerge.

In the current study, we adopted this approach to shed light on the visual experience necessary for creating a human-like, view-invariant representation of faces with DCNNs. The role of experience in human face recognition is well-established. Studies have shown that face recognition is better for familiar than unfamiliar faces (8) and for faces from own race than other race faces for which we have greater visual experience (9–11). Moreover, developmental studies indicate that face recognition gradually improves with development, including the ability to generalize across different images of the same individual (12,13). Many studies have emphasized the importance of experience with variable face images for successful face recognition (12,14–16). However, systematic manipulation of human real-life experience with faces is not possible and it is therefore hard to determine a direct link between the visual diet that humans are exposed to and its contribution to the generation of a face representation that enables their face recognition abilities.

In a recent set of studies, Abudarham and colleagues (6,17,18) discovered that humans are sensitive to a subset of facial features that are critical for face identification. Replacing these features changed the identity of a face (see supplementary Figure 1). Moreover, Abudarham and colleagues (2016) found that human sensitivity to these critical features remained invariant across variations in head pose, which makes them potentially useful for view-invariant identity classification. They further revealed that face-trained, but not object-trained DCNNs, showed similar sensitivity to this subset of facial features. This indicates that experience with faces is necessary to learn to use these features for identity classification. These findings are also consistent with recent studies showing human-like face effects such as the face inversion effect and other-race effect in face-trained but not object-trained DCNNs (19,20). Furthermore, sensitivity to these critical features and the generation of a view-invariant representation were found in higher layers of the face-trained network, whereas earlier layers showed no preference to this subset of face features and evidence for a view-specific face representation (6). This human-like representation enables us to link between humans and DCNNs view-invariant representations and examine the type of visual diet that is required for the development of successful identity classification.

To this end, in the current study we systematically manipulated the amount and type of visual-diet and examined its effect on the generation of a view-invariant representation, sensitivity to human-like critical features and identity classification in DCNNs. In particular, we manipulated experience by gradually increasing the number of within-identity images or between-identity images to examine their relative contributions to the generation of human-like, view-invariant representation. Concurrent emergence of these representations as a function of the visual diet would suggest that they depend on similar features. Data reported in the results section can be found in this OSF link: https://osf.io/huzkp/?view_only=dfc6c75cc12b424d851794be43ca3f44

## Results

To assess the effect of the visual diet on the generation of a view-invariant representation of face identity, we trained a DCNN (VGG-16) with the following training diets: We created 64 subsets of face images, which included all possible combinations of 2, 5, 10, 50, 100, 200, 500, 1000 identities and 1, 5, 10, 20, 50, 100, 200, 300 images per identity. For models that are trained with a relatively smaller number of faces (1-100 identities models), we trained the model with thirty different sets of faces to avoid stimulus specific effects. The results were then averaged across the thirty models for each condition.

### Effects of visual diet on face identity classification

We measured the performance of the DCNNs on a standard face verification task, Labeled Faces in the Wild (LFW) benchmark (21). For each of the networks we extracted the representation in the penultimate layer (FC7) and assessed performance on a same-different identity task (see methods). Figure 1 shows that accuracy improves for DCNNs trained on larger number of images. Both the number of different identities as well as the number of images per identity were needed to improve performance. Accuracy did not exceed 75% if the number of identities was below 10 for any number of images per identity (up to 300 images per identity) or if the number of images per identity was below 5 for any number of identities (up to 1000 identities) (See Supplementary Table 1 for complete report of performance levels). This suggests that identity face classification requires experience with images of different identities but also with different images of the same identity.

**Figure 1:**
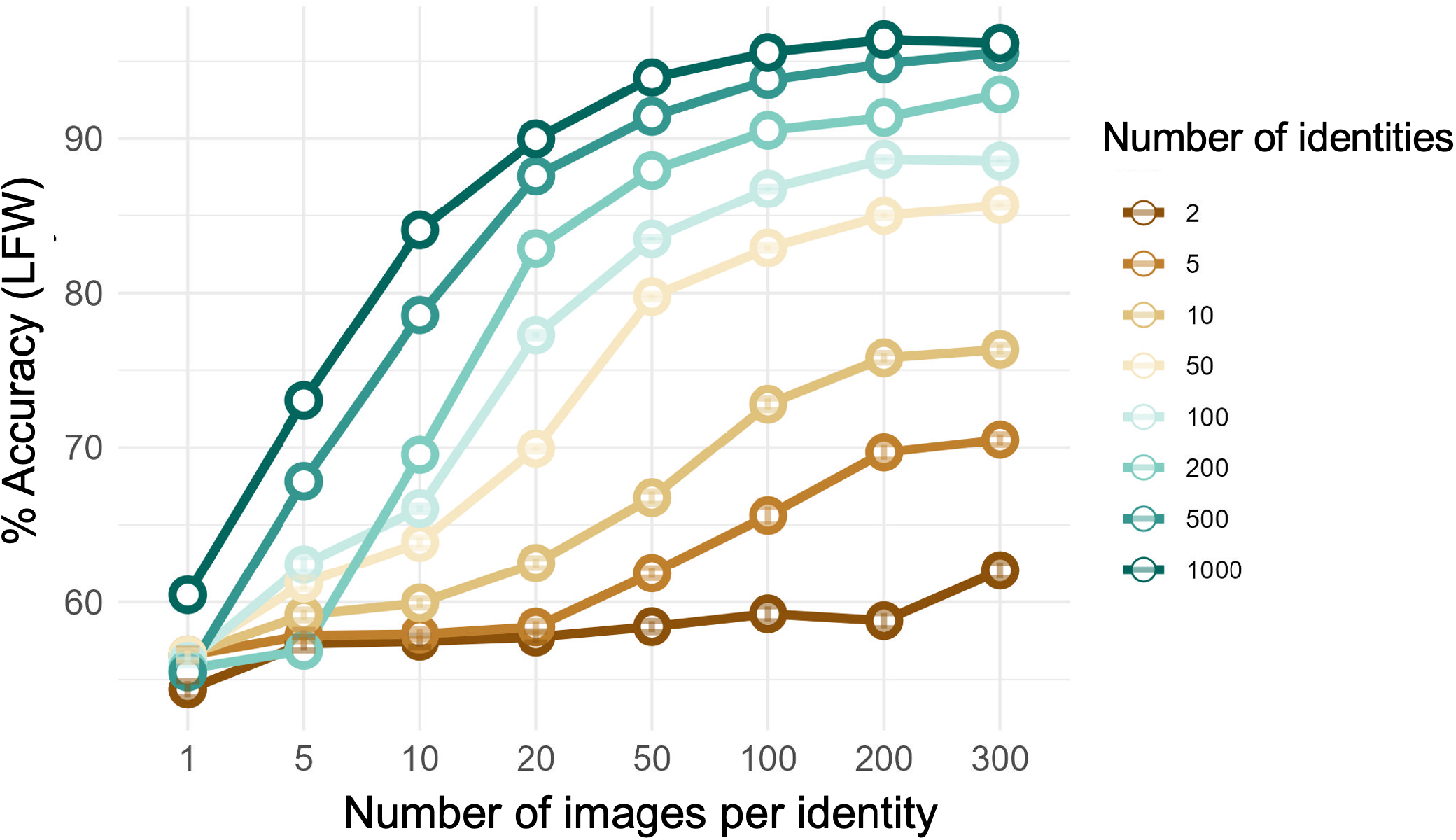
Accuracy on a face verification task with Labeled faces in the wild (LFW) benchmark. Performance gradually improves with increase in the number of identities as well as the number of images per identity.

We next assessed how this experience changes the representation from a view-specific to a view-invariant representation.

### The emergence of a view-invariant representation

To evaluate whether a representation that is generated by a DCNN is view-specific or view-invariant, we used face images of 15 identities from which we generated the following four types of pairs: same identity: same view (frontal), same-identity: different view (frontal vs. quarter view), same-identity: different view (frontal vs. half view), different identity-same view (frontal) (Figure 2A). We measured the Euclidean distance between the feature vectors of the four face pairs for two base-line representations: *Pixel-based representation*, which was the raw pixel values of the test face images. *Identity-based representation*, which was the representation in the penultimate layer of a fully face-trained DCNN (>8000 identities with approximately 300 images per identity, see methods). The distance between the pixel-based representations of each pair of faces showed a view-specific representation as indicated by a larger distance between same identity-different view face pairs (light blue bars) than between different identity-same view face pairs (red bar) (Figure 2B). The distance between the representations of each pair of faces based on the penultimate layer of the fully-trained network revealed a view-invariant representation as manifested by a larger distance between different identity-same view face pairs than between same identity-different view face pairs (Figure 2C) (6).

**Figure 2:**
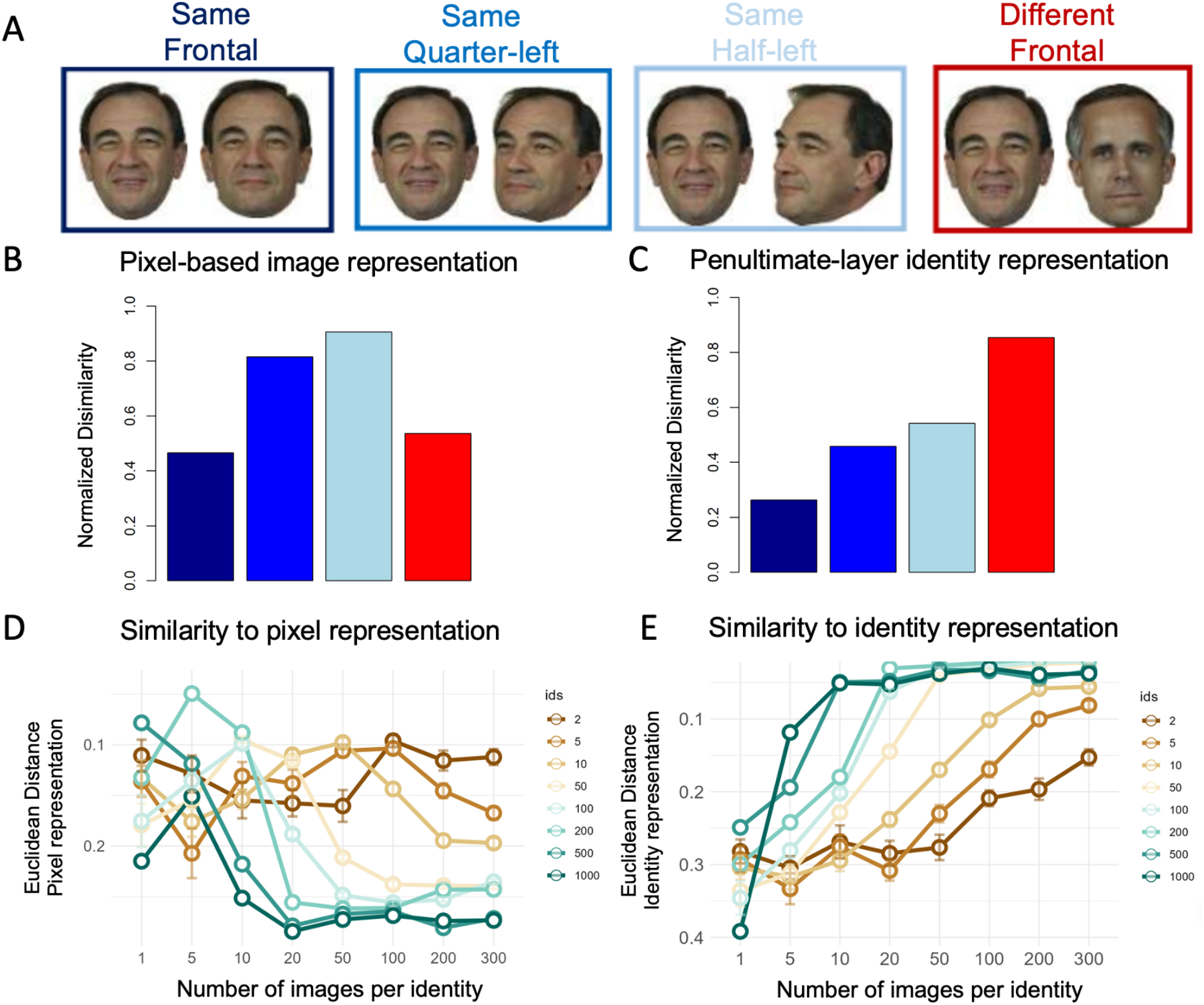
The emergence of a view-invariant face representation with visual experience. A. The four types of face pairs used to test a view-invariant representation. Same identity same view (dark blue), same identity quarter-left (light blue), same identity half-left (lightest blue) and different identity same view (red). B. Disimilarity scores (normalized distance) between the four types of face pairs based on the pixel layer reveals a view-specific representation. C: Disimilarity scores (normalized distance) between the four types of face pairs based on the penultimate layer of a fully face-trained DCNN reveales a view invariant representation. D. The similarity (measured by Euclidean distance) of each DCNN with the pixel-based representations (panel B) is higher for DCNNs trained on smaller number of images. E. The similarity (measured by Euclidean distance) of each DCNN with the identity-based representation (panel C) is higher for DCNNs trained on larger number of images. To see the representations of each DCNN see Supplementary Figure 3.

Next, we measured the similarity between the representations generated for the same four face pairs in each of the trained models with the pixel-based and the identity-based representations. We did that by calculating the Euclidean distance between the distribution of the four types of face pairs of the *pixel-based* representation (Figure 2B) and *identity-based* representation (Figure 2C) with the distribution of the four types of face pairs in each of the 64 DCNN trained models (see supplementary Figure 2). The similarity to the *pixel-based* distribution is presented in Figure 2D and to the *identity-based* distribution in Figure 2E. We found that DCNNs that are trained on smaller number of identities and images per identity generate representations that are more similar to a view-specific, image-based representation, and DCNNs that are trained on larger number of identities and images per identity generate a representation that is more similar to a view-invariant, identity-based representation. In particular, we see that a DCNN that is trained on large number of identities (500 or 1000) generate a view-invariant representation with only 10 images per identity.

### Sensitivity for human-like critical features

To evaluate whether the representations of DCNNs are sensitive to human-like view invariant critical features, we measured the distance between representations of four types of face pairs of 25 different identities (not included in the train set). Figure 3A shows an example of each type of face pairs: “Same identity” are different images of the same identity, “Non-critical features” are same identity face pairs in which non-critical features were replaced (see supplementary Figure 1 bottom); “Critical features” are same identity face pairs in which critical features were replaced (see supplementary Figure 1 top); “Different identity” face pairs. Figure 3B shows the Euclidean distances between these face pairs based on their pixel-based representations. The distance was similar for faces that differ in critical or in non-critical features, indicating that pixel information is not sensitive to human-like critical features more than non-critical features. Figure 3C shows the Euclidean distances between representations of the same face pairs, based on the penultimate layer of a fully face-trained DCNN (6,17). Here we see a much larger distance between faces that differ in critical features than faces that differ in non-critical features, indicating that the identity-based representation is sensitive to human-like critical features. We also show that faces that differ in critical features are as different as different identity faces, indicating that changing them is similar to changing the identity of a face.

**Figure 3:**
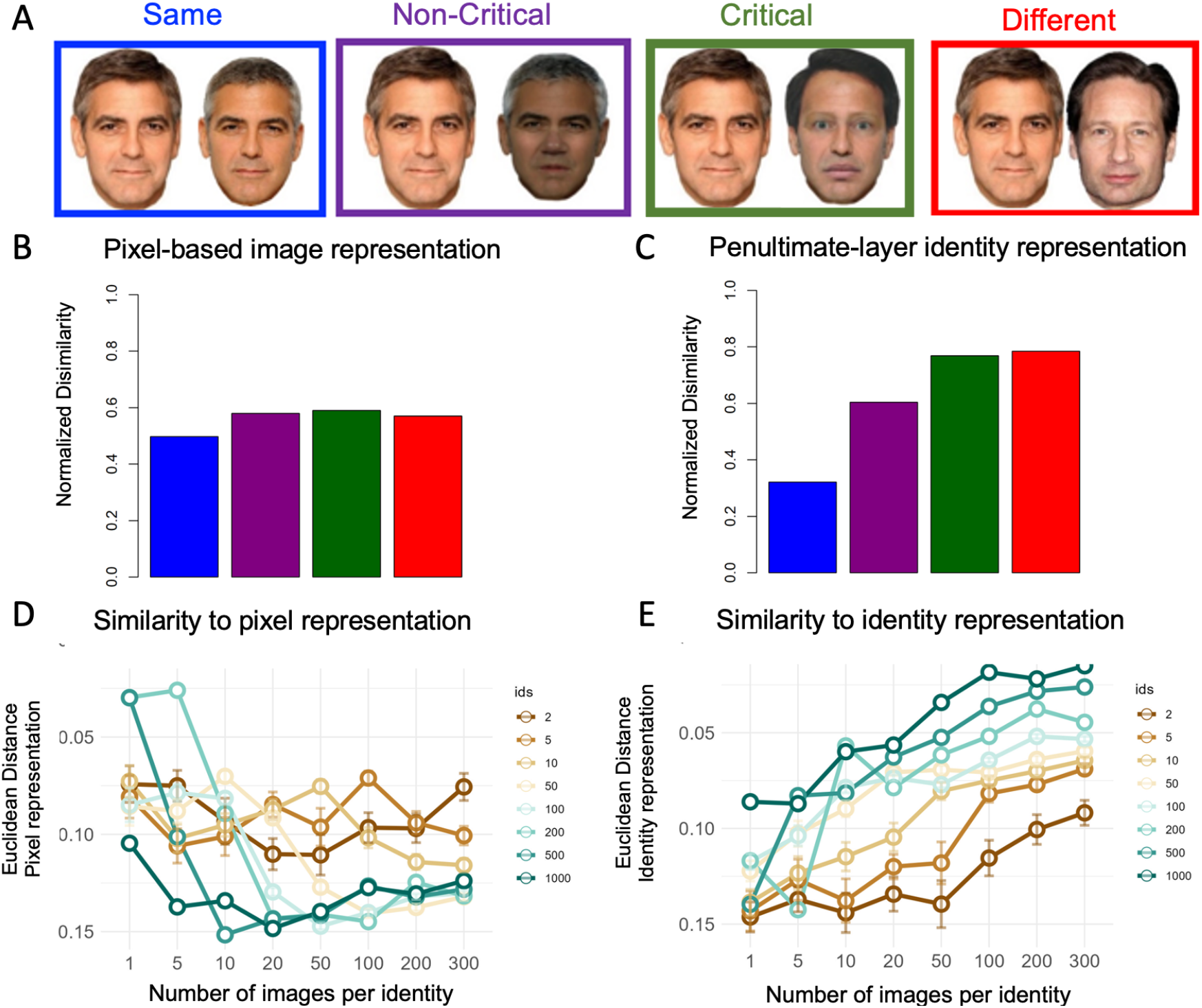
The emergence of sensitivity to critical features: A. The four types of face pairs that are used to test sensitivity for critical features. Same identity (blue), non-critical features changed (purple), critical features changed (green) and different identity (red). B. Disimilarity scores (normalized distance) between the four types of face pairs based on the pixel layer reveals no sensitivity to human-like critical features. C: Disimilarity scores (normalized distance) between the four types of face pairs based on the penultimate layer of a fully face-trained DCNN reveales high sensitivity to human-like critical features. D. The similarity (measured by Euclidean distance) of each DCNN to the pixel-based representations (panel B) is higher for DCNNs trained on smaller number of images. E. The similarity (measured by Euclidean distance) of each DCNN to the identity-based representation (panel C) is higher for DCNNs trained on larger number of images. To see the representations of each DCNN see Supplementary Figure 3.

Next, we measured the similarity of the representations to the four face pairs that were generated by each of the face-trained models with the pixel-based and identity-based representations. We calculated the Euclidean distance between the distribution of the four types of face pairs of the *pixel-based* representation (Figure 3B) and *identity-based* representation (Figure 3C) with the distribution of the four types of face pairs in each of the 64 DCNN trained models (see supplementary Figure 3). The distances from the *pixel-based* distribution are presented in Figure 3D and from the *identity-based* distribution in Figure 3E. DCNNs that were trained on smaller number of images were more similar to the image-based representation showing no sensitivity to critical features over non-critical features. Whereas DCNNs that were trained on a larger number of images were more similar to models that are sensitive to human-like critical features.

Abudarham and Yovel (18) suggested that humans are sensitive to critical features because they enable a view-invariant representation of face identity which is needed for successful face recognition across different appearances of the same identity. To examine these correspondences, we computed the correlations between the distance between pairs of faces of same identity that differ in non-critical featrures (Figure 4A,B) and the distance between pairs of same identity faces that differ in critical features (Figure 4C,D) and examined their correlations across all 64 DCNNs with accuracy on face verification based on the LFW benchmark (Figure 4A,C) and with a measure that indicates a view invariant representation (Figure 4B,D). The view invariant measure was the difference between different face pairs (red bar in Figure 2) and same identity different head view face pair (light blue in Figure 2). As can be seen in Figure 2, the relative distance between these two bars changes drastically between the pixel-based representation in which different identity-same view faces are more similar than same identity-different view faces (Figure 2B) and the identity-based, view-invariant representation in which the distance between same identity – different view faces is much smaller than same view different identity faces (Figure 2C). Thus, a positive difference between these face pairs indicate a view invaraint represnetaion and a negative distance a view-specific representation).

**Figure 4:**
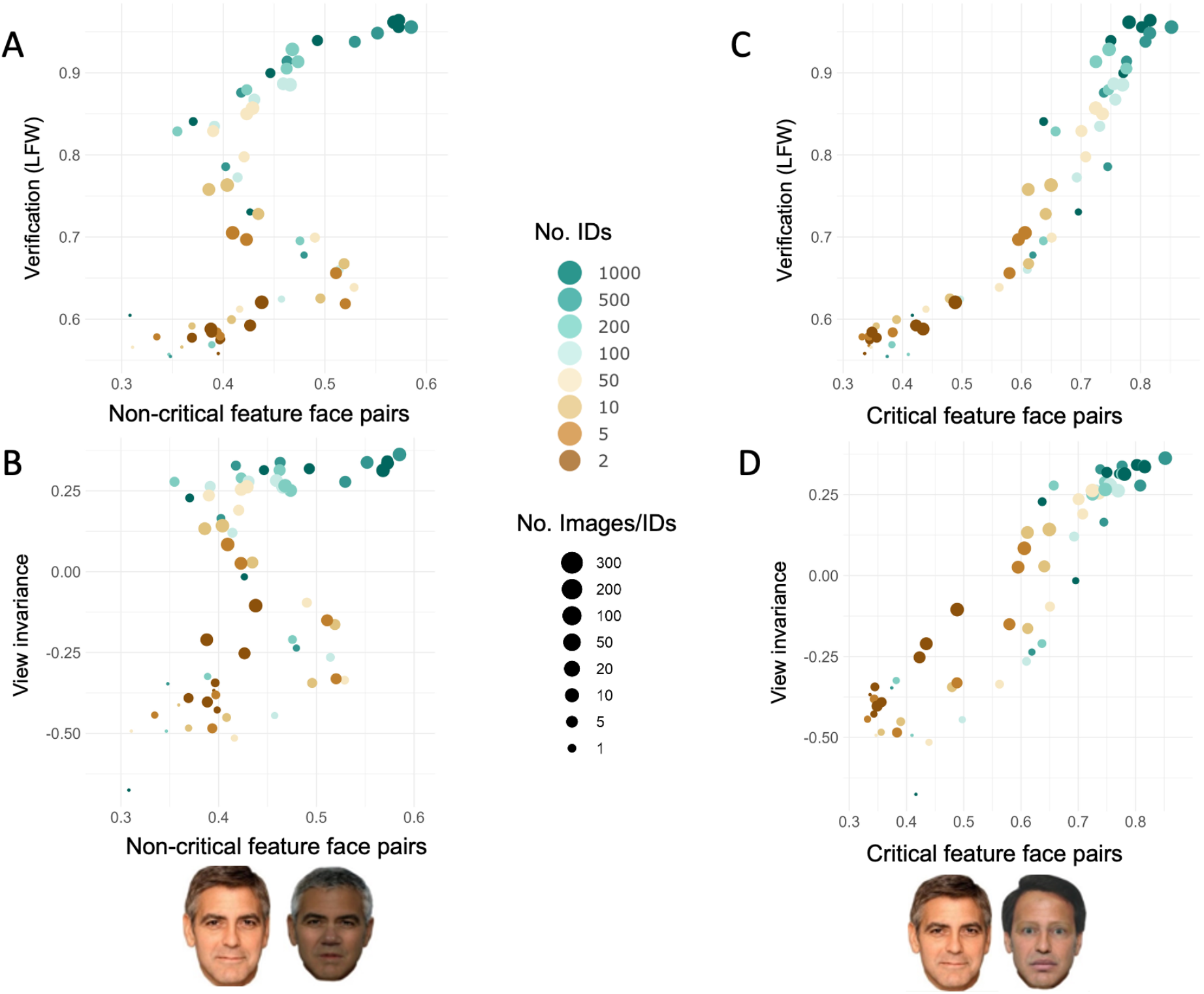
Sensitivity to human-like critical features is correlated with performance on face verification and the emergence of view invariance representation: A. Sensitivity to non-critical features, measured by the distance between same identity faces that differ in non-critical features (see supp figure 1), and performance on face verification task do not emerge concurrently as a function of experience. B. Sensitivity to non-critical features do not emerge concurrently with a view-invariant representation as a function of experience. C. Sensitivity to critical features, measured by the distance between same identity faces that differ in critical features (see supp figure 1), and performance on face verification task emerges concurrently as a function of experience. D. Sensitivity to critical features emerges concurrently with a view-invariant representation as a function of experience.

Figure 4 shows a strong linear relationship between sensitivity to critical featurs (distance between pairs of faces that differ in critical features) and accuracy of the DCNN on the LFW benchmark (Figure 4C, r(62) = 0.95). It also shows a strong linear relationship between sensitivity to critical features and the emergence of a view-invariant representation (Figure 4D, r(62) = 0.93). There was no such linear relationship between sensitivity to non-critical features (distance between pairs of faces that differ in non-critical features) and performance on face verification task (Figure 4A, r(62) = 0.56) or the emergence of a view invariant representation (Figure 4B, r(62) = 0.47).

## Discussion

Successful face recognition depends on the ability to generalize across different images of the same identity and discriminate between images of different identities. The goal of the current study was to leverage the success of DCNNs in face recognition and their similarity to human-like representations (6,19,22), to examine whether success on a face verification task, the emergence of a view-invariant representation and sensitivity to human-like critical facial features emerge concurrently as a function of the amount and type of experience with faces. Our findings show that increasing both the number of images per identity and number of identities, concurrently improved verification accuracy, the emergence of a view-invariant representation and sensitivity to human-like critical facial features. These findings suggest a critical role for experience with faces in the generation of these representations.

For many years cognitive scientists and computer scientists have attempted to reveal the critical features that enable human-level face recognition performance. Despite the success of current machine learning algorithms to recognize faces at, or even above, human-level performance, it is still unknown which features are used by these algorithms to perform this task. Studies in humans revealed a subset of facial features for which humans showed high perceptual sensitivity. Furthermore, changing these features changed the identity of the face (see Supplementary Figure 1) indicating their importance for human face recognition (17). Abudarham and colleagues (2016) further suggested that these features enable a view-invariant representation, as they remain invariant across different head-views (18). In the current study, we were able to link these two phenomena and their relationship with verification accuracy by showing that they emerge concurrently as a function of the amount of experience with faces during training (Figure 4). These findings suggest that these identity-based representations rely on similar features.

The relevance of these findings to human face recognition should be evaluated considering the nature of human experience with faces during development. Recent studies that have used head-mounted cameras on infants’ foreheads during the first year of their life show that during this period, they were primarily exposed to three main identities from myriad of different appearances and head-view (23). It is only later during development that the number of identities start increasing reaching a few thousands of familiar identities in adult. (24) Indeed, performance in face recognition improves slowly and requires several years to reach adult level performance (13). To better learn about effects of human-like experience from face recognition algorithms, it is necessary to train the algorithms on a more human-like type of experience with faces, which is different from the training set and training protocols of current face recognition algorithms (25,26). Another important difference between human and face recognition algorithms is that human face recognition primarily concerns the recognition of familiar faces (8,27), whereas face recognition algorithms are trained to achieve impeccable classification of unfamiliar identities. In the current study, we used unfamiliar faces to test the representation of face recognition algorithms and learn about their ability to generalize to unlearned examples. However, if the goal of the human face recognition system is to only classify socially relevant familiar identities, computer algorithms that aim to model human face recognition should take this consideration into account. Given that an important aspect of familiar face recognition is their semantic representations, the development of familiar face recognition may be better modelled with multi-modal visual-semantic algorithms (26,28).

Recent studies that examined human-like representations in DCNNs revealed other similarities between humans and DCNNs. This includes a much larger drop in performance for inverted than upright faces than the drop that is found for objects (19,20). The Thatcher illusion in which distorted faces look more similar to normal faces when they are inverted than upright was also found in face-trained but not object-trained DCNNs (29) A drop in performance for the race of faces that the algorithms was not trained on (i.e. lower performance for Asian faces in a DCNN trained on White faces) is also typically found in DCNNs similar to the human other race effect (7,19). The approach that we used in the current study enables us now to ask what kind of experience is required for these human-like representations to emerge.

In summary, recent advances in machine learning that enable face recognition algorithms to reach human-level performance, and the similarity between the representations generated by humans and machines (6,19,29), offer us new computational tools to explore the factors that mediate human face recognition. Future studies will further investigate the contribution of more specific characteristics of face images, such as their pose, expression, and lighting, to the generation of a view-invariant representation and sensitivity to view-invariant human-like critical features.

## General Methods

### Model

We used VGG-16 (3) as a the base model, which we trained on different numbers of face images. We selected this model because it has been often used in previous studies (6,19,22). The representations used in the study are extracted from the penultimate layer (FC7).

### Train Dataset

We used the VGGFace2 dataset (30) to train our networks. VGGFace2 is a large-scale face recognition dataset developed by the Visual Geometry Group at the University of Oxford. It contains over 3 million images of more than 9,000 individuals, with each individual represented by several hundred images. The images were collected from a variety of sources and were annotated with bounding boxes and labels indicating the identity of the individuals.

### Training protocol

We created 64 subsets of face images, which included all possible combinations of 2, 5, 10, 50, 100, 200, 500, 1000 identities and 1, 5, 10, 20, 50, 100, 200, 300 images per identity. For the small training sets (1-100 identities, with all possible images per identity), we trained each DCNN on thirty different data sets to obtain robust performance measure of their representations/performance. The results were then averaged across the thirty networks. Representations were extracted also from the fully-trained model that was trained on the whole VGGface2 data set.

### Stimuli

#### View-specific and view-invariant representations

To examine whether the network generates a view-specific or a view-invariant representation, we used images of 15 identities from the color FERET face-image dataset. For each identity, we selected four images: a “reference” frontal image, a second “frontal” image that is different from the reference image, a quarter-left image, and a half-left image. All images were of adult Caucasian males, well-lit, with no glasses, hats or facial hair. The images were cropped just below the chin to include only the face, including the hair and ears. This resulted in four types of face pairs: “Same-Frontal”, “Same-quarter view”, “Same-half view” and “Different-Frontal” (See Figure 2A for examples of the four types of face pairs).

#### Critical features for face recognition

We used 25 face identities to generate image pairs. For each of the 25 identities, we used an original image, an image with modified critical features, and an image with modified non-critical features (for more information about how the face images were created see (18). We also used a different unmodified image of the same person, which we used as a reference image. This allowed us to create four image pairs: the “Same” pair, which compares the reference image to the original image, the “Different” pair, which compares the reference image to a reference image of a different identity, the “Critical features” pair, which compares the reference image to the original image with different critical features, and the “Non-critical features” pair, which compares the reference image to the original image with different non-critical features. (See Figure 3A for examples of the 4 types of face pairs).

### Stimuli

#### Performance measures

We measured the performance of the trained DCNNs on a face verification task using the standard Labeled Faces in the Wild (LFW) benchmark (21) including 6,000 pairs of face images for testing. These pairs consist of positive pairs, where both images show the same person, and negative pairs, where the two images show different people. The goal of the face verification task is to determine if the two images in each pair belong to the same person or not. We assessed the models’ performance by measuring the cosine distance between the embeddings of pairs of faces. If the distance was smaller then a pre-determined threshold, the images were classified as the same person, otherwise they were classified as different. The accuracy values reported here reflect performance achieved using the optimal threshold for each model.

#### Quantifying view-invariance of face-representations in DCNNs

We calculated the Euclidean distances between the penultimate layer (fc7) embeddings of the following pairs of faces: same identity faces - same view, same identity faces - quarter view, same identity faces - half view, and different identity faces – same view (as shown in Figure 2A) for 15 different identities. (The face alignment procedure failed to detect 4 of the half-view faces, so we only had 11 face pairs in the frontal-half-view condition). The distance scores were normalized by dividing the measured distances by the maximal distance value in each run across all stimuli and conditions. This resulted in a normalized score that ranged from 0-1. These distance scores were calculated for each of the 64 DCNNs (see Supplementary Figure 2), as well as for a *pixel-based representation* based on the pixel values of the images and for the *identity-based representation* based on the values of the penultimate (last hidden – fc7) layer of the fully face-trained DCNN (6). Finally, to measure the similarity between each of the 64 trained DCNNs and the baseline models (*pixel-based* and *identity-based*), we calculated the Euclidian distance between the normalized mean distances of the four face pairs (dividing the distance of each pair by the sum of the distances of the four pairs) of each trained DCNN with each baseline models. Smaller distances indicate that the DCNN is more similar to the pixel model (Figure 2D) or the identity model (Figure 2E).

#### Measurement of sensitivity to critical features

We calculated the Euclidean distances between the representations of the following four conditions: Same, Non-Critical, Critical and Different (See Figure 3A). Each condition includes 25 image pairs. Distances were calculated for a *pixel-based representation* based on the pixel values of the images and for the other representations based on the penultimate layer. We then performed the same analysis that is described in the previous section to measure the similarity of each of the 64 models with a pixel-based or an identity-based representations (see Supplementary Figure 3).

## Acknowledgement

This work was funded by an ISF grant 917/21 to GY

## Supplementary Material

**Supplementary Figure 1:**
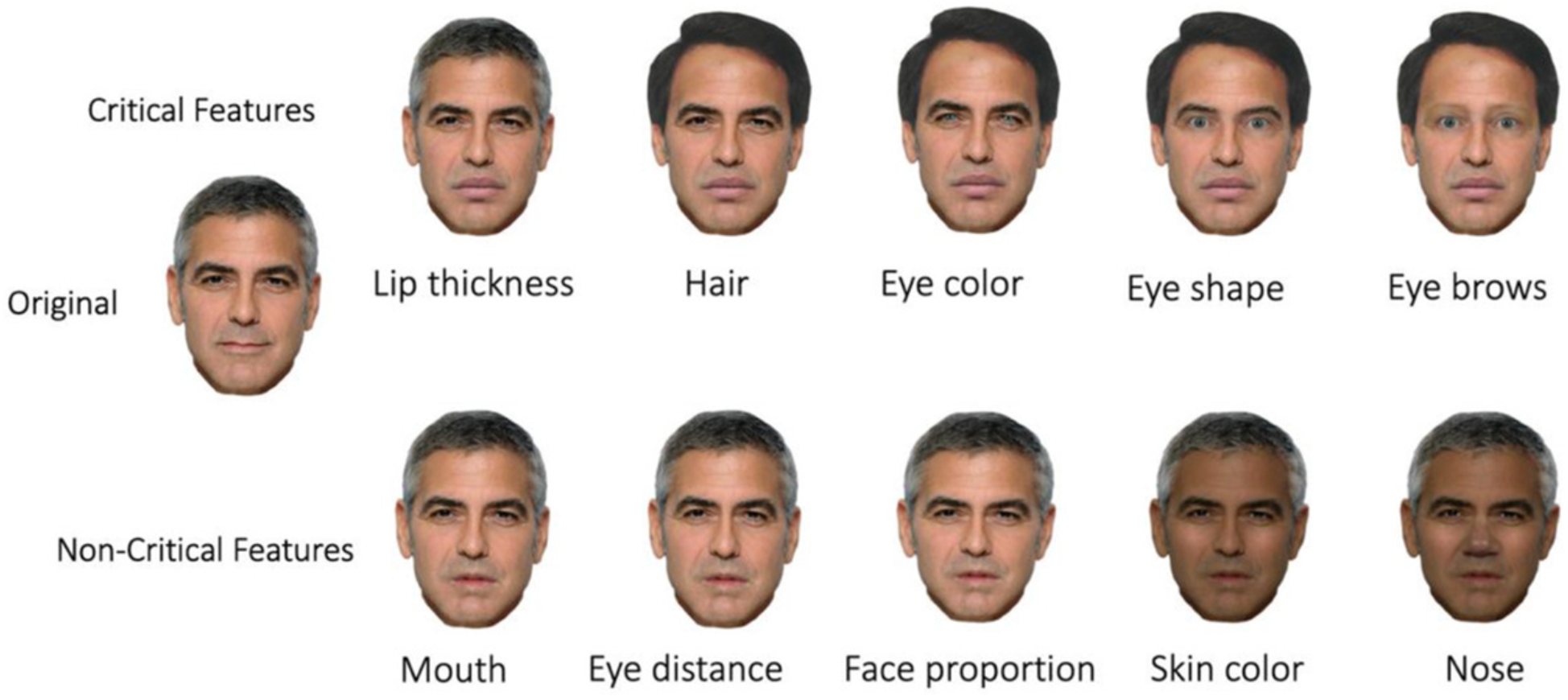
An example of the effect of changing critical or non-critical features on the face of George Clooney. Upper row: Changing five critical features gradually. Bottom row: Changing five non-critical features gradually. Abudarham et al (2019) shows that changing five critical features changes the identity of the face, whereas changing five non-critical features does not change the identity of a face.

**Supplementary Figure 2:**
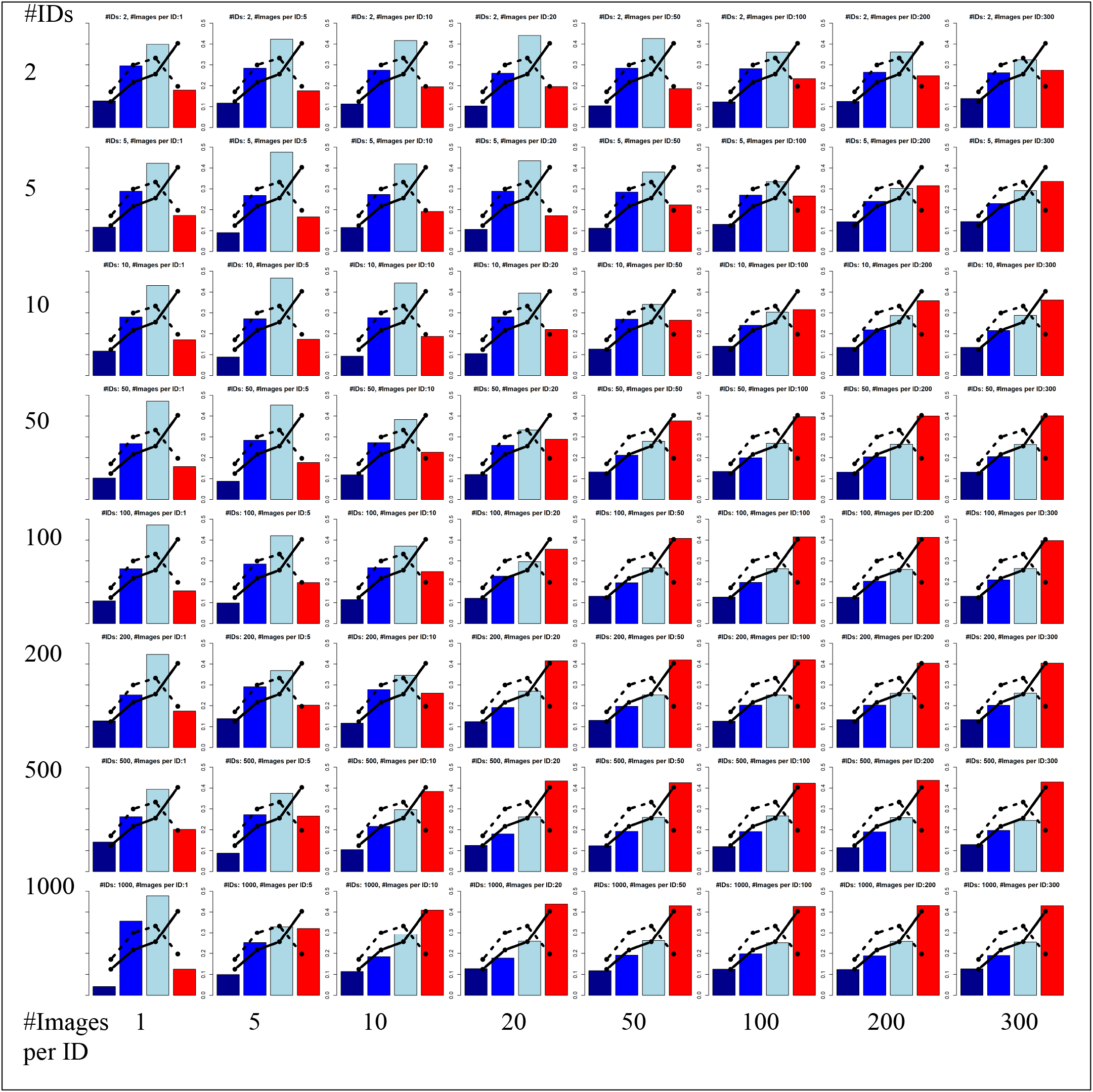
A view invariant representation. The Euclidean distance between the face pairs shown in Figure 2A for each of the 64 DCNNs that were trained on different number of identities and different number of images per identity. The lines indicate the two baseline models: the pixel-based representations (dotted line, see Figure 2B) and the identity-based representation (solid line, see Figure 2C). Figure 2D,E show the similarity between the pattern of findings in each DCNN with the baseline models.

**Supplementary Figure 3:**
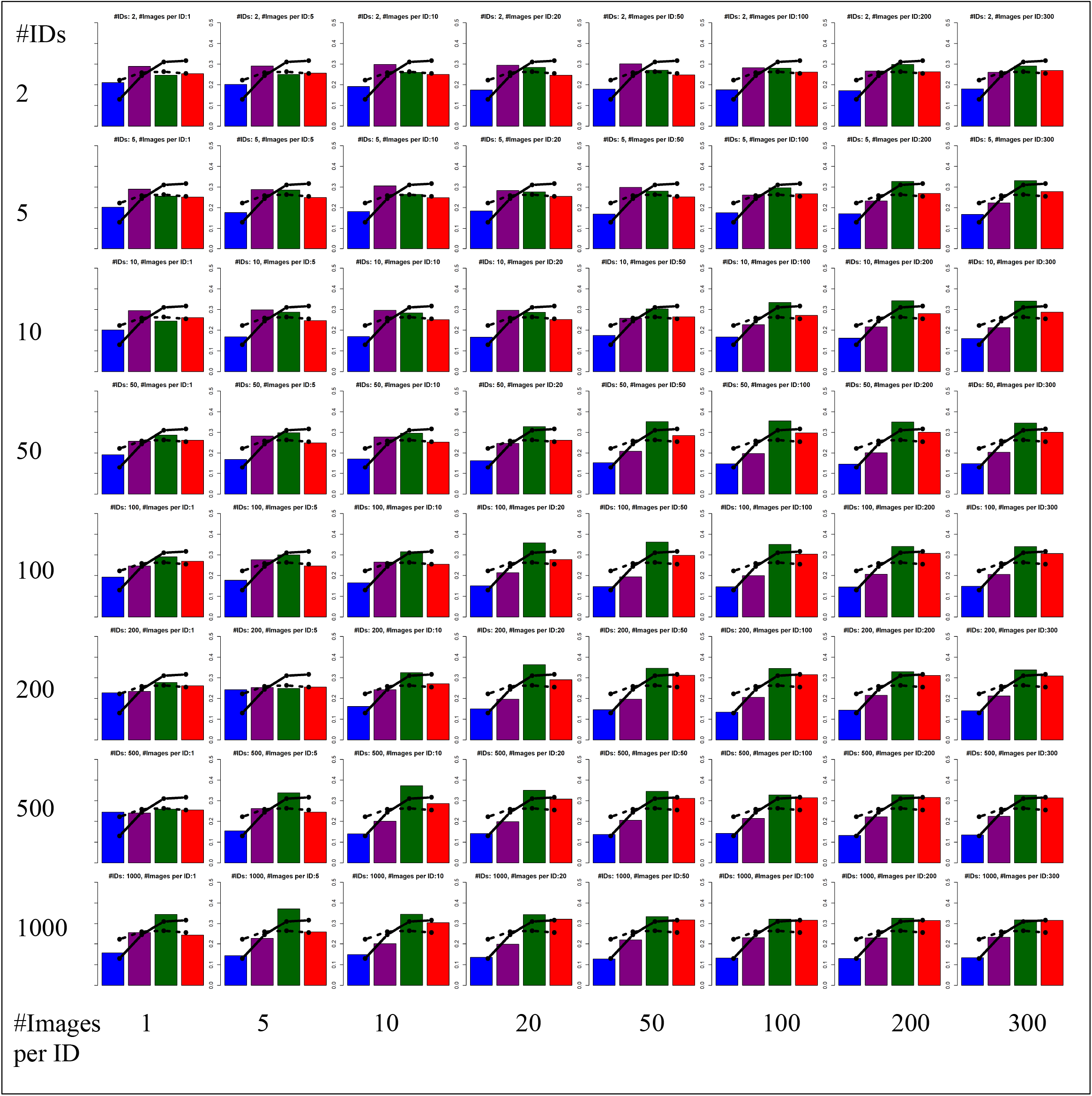
Sensitivity to critical features. The Euclidean distance between the face pairs shown in Figure 3A for each of the 64 DCNNs that were trained on different number of identities and different number of images per identity. The lines indicate the two baseline models: the pixel-based representations (dotted line, see Figure 3B) and the identity-based representation (solid line, see Figure 3C). Figure 3D,E show the similarity between the pattern of findings in each DCNN with the baseline models.

**Supplementary Table 1:** Accuracy on the face verification task for each of the DCNNs that were trained on different number of identities (IDs) / different number of images per identity (see Figure 1).

| IDs/Images | **1** | **5** | **10** | **20** | **50** | **100** | **200** | **300** |
| --- | --- | --- | --- | --- | --- | --- | --- | --- |
| **2** | 0.56 | 0.58 | 0.57 | 0.58 | 0.58 | 0.59 | 0.59 | 0.62 |
| **5** | 0.57 | 0.58 | 0.58 | 0.58 | 0.62 | 0.66 | 0.70 | 0.71 |
| **10** | 0.57 | 0.59 | 0.60 | 0.63 | 0.67 | 0.73 | 0.76 | 0.76 |
| **50** | 0.57 | 0.61 | 0.64 | 0.70 | 0.80 | 0.83 | 0.85 | 0.86 |
| **100** | 0.56 | 0.62 | 0.66 | 0.77 | 0.83 | 0.87 | 0.89 | 0.89 |
| **200** | 0.56 | 0.57 | 0.70 | 0.83 | 0.88 | 0.91 | 0.91 | 0.93 |
| **500** | 0.55 | 0.68 | 0.79 | 0.88 | 0.91 | 0.94 | 0.95 | 0.96 |
| **1000** | 0.60 | 0.73 | 0.84 | 0.90 | 0.94 | 0.96 | 0.96 | 0.96 |

